# Enhancing cellular production of fucoxanthin through machine learning assisted predictive approach in *Isochrysis galbana*

**DOI:** 10.1101/2024.04.26.591373

**Authors:** Janani Manochkumar, Annapurna Jonnalagadda, Aswani Kumar Cherukuri, Brigitte Vannier, Dao Janjaroen, Siva Ramamoorthy

**Affiliations:** School of Bio Sciences and Technology, Vellore Institute of Technology, Vellore 632014, India; School of Computer Science & Engineering, Vellore Institute of Technology, Vellore 632014, India; School of Computer Science Engineering and Information Systems, Vellore Institute of Technology, Vellore 632014, India; Cell communications and Microenvironment of Tumors Laboratory, UR 24344, University of Poitiers, France; Department of Environmental Engineering, Faculty of Engineering, Chulalongkorn University, Bangkok 10330, Thailand

**Author notes:** **Corresponding author** Siva Ramamoorthy, School of Bio Sciences and Technology, Vellore Institute of Technology, Vellore 632014, Tamil Nadu, India **Email:**.

**Keywords:** Fucoxanthin, *Isochrysis galbana*, Phytohormones, Machine Learning, Prediction

## Abstract

The marine microalgae, *Isochrysis galbana* is a prolific producer of fucoxanthin which is a xanthophyll carotenoid with substantial global market value boasting extensive applications in the food, nutraceutical, pharmaceutical, and cosmetic industries. Although supplementation of different phytohormones to medium enhances fucoxanthin production, the quantification of pigment by conventional means is time-consuming and labor-intensive. This study addressed the multiple methodological limitations of HPLC-based fucoxanthin quantification and emphasized the need to develop a Machine Learning (ML) model as optimization and precise prediction remain a challenging task. Hence, an integrated experimental approach coupled with ML models was employed to predict fucoxanthin yield by supplementation of various phytohormones. The accuracy of fucoxanthin prediction excluding and including hormone descriptors was compared and evaluated using the ML models namely Random Forest (RF), Linear Regression (LR), Artificial Neural Network (ANN), and Support Vector Machine (SVM). RF model provided the most accurate prediction excluding hormone descriptors with coefficient of determination (*R*^2^ = 0.809) and root-mean-square error (*RMSE* = 0.776) followed by the ANN model with (*R*^2^ = 0.722) and (*RMSE* = 0.937). The inclusion of hormone descriptors for training and pre-processing of data further improved the fucoxanthin prediction accuracy of the RF model to (*R*^2^ = 0.839) and (*RMSE* = 0.712) and ANN model to (*R*^2^ = 0.738) and (*RMSE* = 0.909). These results indicated that the combination of low-cost, Ultraviolet (UV) spectrometric-based fucoxanthin quantification coupled with ML algorithms can be efficiently used for reliable prediction and enhanced fucoxanthin production, therefore highlighting a promising approach and furnishing invaluable insights towards the commercialization of microalgal fucoxanthin production.

## Introduction

Microalgae are a diverse group of photoautotrophic organisms with a promising source of bioactive compounds including polysaccharides, fatty acids, carotenoids, phytosterols, and phenols with beneficial applications (Lopes et al. 2021). Recently, extensive research has contributed to investigating the potential of microalgae to synthesize value-added metabolites owing to its simple cell organization, enhanced accumulation of lipids, rapid life cycle, steady growth rate, non-toxic, biodegradable and utilization of CO_2_ as a carbon source for growth (Chong et al. 2023). Marine pigments thus have evolved to be an effective alternative in food, therapeutic, and cosmetic applications (Manochkumar et al. 2022).

Fucoxanthin is a xanthophyll marine carotenoid that is abundantly found in the thylakoid membrane of chloroplasts in macroalgae and its distribution varies within chloroplast in microalgae (Foo et al. 2021; Manochkumar et al. 2022). Among all the carotenoids, fucoxanthin profusely contributes to more than 10% of estimated total carotenoid production globally (Lopes et al. 2021). Currently, the global market value of fucoxanthin upholds an average annual growth rate of 5% representing an increase from 199.48 million USD in 2022 to 280.7 million USD in 2029 (Market Reports World, 2023). Due to its enormous applications and valuable bioactivities, the cost of purified fucoxanthin ranges from 40,000 to 80,000 USD/kg depending on the concentration and purity of the compound (Khaw et al. 2022). Despite its promising applications, fucoxanthin remains limited in availability as it has not yet been fully commercialized. Indeed, synthetic production of fucoxanthin is not feasible due to its complexity, hence microalgae were explored as an effective and reliable source for the fucoxanthin production (Lopes et al. 2021). It is evident that fucoxanthin plays a significant role in microalgae by absorption of photons thus regulating photosynthesis and aiding photoprotection to chlorophyll from photodamage (Miyashita et al. 2020)

*Isochrysis galbana* belongs to the class of flagellated marine microalgae that shows a higher accumulation of lipids, omega-3 polyunsaturated fatty acids, and fucoxanthin. The absence of cell wall in this species allows for an easy extraction of fucoxanthin during downstream processing (Sun et al. 2019). The inherent capability of microalgae to produce a higher fucoxanthin yield, short life cycle, independent of seasonal variations, could be cultivated all year round, and not competing for land makes it a promising source of fucoxanthin (Yusof et al. 2022).

The common method used to estimate the pigment concentration is high-performance liquid chromatography (HPLC) which requires long extraction and column time for each run, is time and cost-consuming, and requires highly skilled persons to maintain the equipment (Chong et al. 2023). Hence, a high throughput method should be simple, accurate, and reliable for the extraction and detection of pigment. Here we employed the equation derived by Wang et al. (2018), for UV spectrometry-based quantification of fucoxanthin. While the HPLC method requires at least 3 hours to quantify the fucoxanthin, this method could detect the fucoxanthin within 5 minutes (Wang et al. 2018). However, this method will not be feasible for large-scale extraction of fucoxanthin due to limited technological advancement (Tang et al. 2023).

Thus, ML models were implemented for the prediction of fucoxanthin yield as it requires less solvent, analysis time and low cost, high accuracy, and good prediction. The accuracy of the prediction of ML models depends on the input variables and the training dataset. In this study, the experimental data of *Isochrysis galbana* supplemented with different types and concentrations of phytohormones was subjected to statistical analysis followed by data preprocessing to train the ML models for fucoxanthin prediction.

The advancements in ML and Artificial intelligence (AI) algorithms have profoundly contributed to the easy search for novel natural product-based drug discovery in the 21^st^ century (Manochkumar and Ramamoorthy 2024). Recently, numerous omics-related datasets have been developed for diverse species of marine organisms and the need to develop and integrate ML algorithms for multi-omics studies has been extensively reviewed (Manochkumar et al. 2023). In crop breeding research, multimodal data from three sensors coupled with ML algorithms were efficiently used in a study for the estimation of the crop harvest index of Faba bean and Pea (Ji et al. 2024). Similarly, ML-based phenotyping combined with optical tomography was used to measure the stomatal density and improve the water use efficiency of Sorghum crop (Ferguson et al., 2021).

In previous studies related to microalgae, an ML model was incorporated to derive a spectrophotometric equation for simultaneously quantifying the concentration of chlorophyll, violaxanthin, zeaxanthin, and lutein from *Chlorella vulgaris* and *Scenedesmus almeriensis* (Victor et al. 2023). Similarly, the convolutional neural network (CNN) model was used to predict the microalgal pigments including chlorophyll-a, phycocyanin, lutein, fucoxanthin, and zeaxanthin from diatoms using experimental data obtained from water samples (Pyo et al. 2022). A hybrid ML-based approach was developed to optimize the production of biomass and phycobiliproteins in *Nostoc* sp. (Saini et al. 2021). Furthermore, Tang et al. (2023) compared linear regression with the ANN model to predict the chlorophyll concentration in *Desmodesmus* sp. and *Scenedesmus* sp. based on RGB, CYMK, and HSL color models. In this study, we constructed two ML frameworks to compare and evaluate the predictive performance of four models on fucoxanthin production from *Isochrysis galbana* by altering the input data parameters. The overall process workflow of experimental and ML setup for fucoxanthin production is depicted in **Figure 1**.

**Figure 1.**
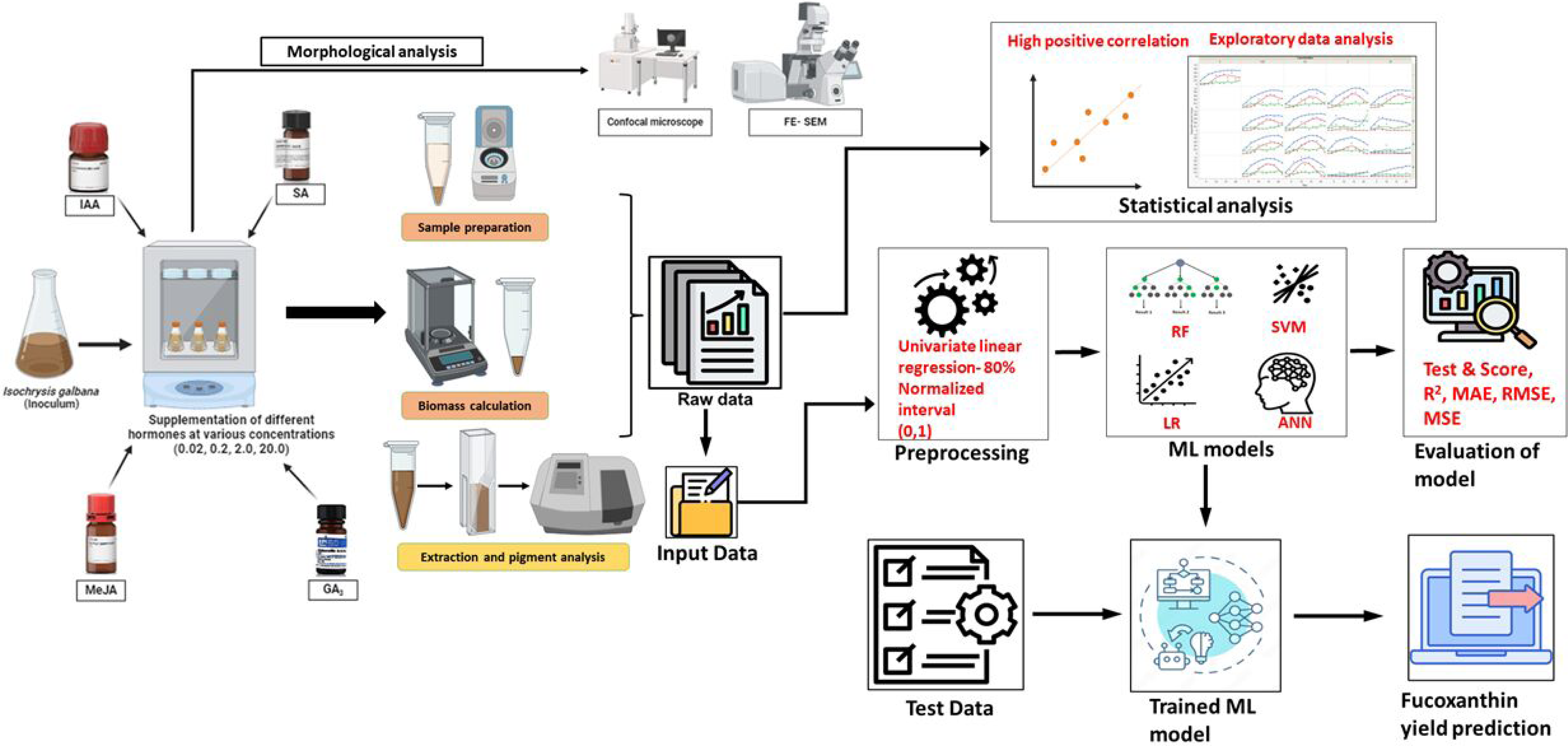
Overall process workflow of experimental and machine learning setup. *Isochrysis galbana* was scaled up and the inoculum was added to medium supplemented with varying concentrations of phytohormones. The growth rate, dry and fresh weight of biomass and fucoxanthin yield were measured on alternate days. Morphological analysis of microalgal cells (exponential phase) was observed using FESEM and confocal microscopic analysis. On the other hand, the raw experimental data was subjected to statistical analysis to understand the characteristic pattern of data. Further, the data is fed as raw data as well as pre-processed data for training the ML models and the performance will be evaluated. Finally, the test data will be fed into the trained ML model to evaluate the prediction of fucoxanthin yield.

## Materials and Methods

### Experimental design

*Isochrysis galbana*, a marine water microalga was obtained from the National Repository for Microalgae and Cyanobacteria, Bharathidasan University. The seawater used for media preparation was collected freshly from Mandapam, Tamil Nadu (9^◦^16′17.9′′N79^◦^07′49.4′′E) and filtered through 0.22 μm membrane filtration system followed by the sterilization using autoclave for 20 min at 121 °C. The salinity and pH of the sterilized seawater must be within 27 ± 1 and 8 ± 0.5 respectively. The experimental setup was maintained under controlled laboratory conditions with optimum temperature (23 ± 2 °C), light intensity (2000 lx), and photoperiod (16 h Dark: 8 h Light) for 30 days.

The *Isochrysis galbana* stock solution was maintained in Conway’s medium for 14 days and its density was adjusted to 2.5 mg/ml of wet biomass using sterile seawater. The algal suspension was then partitioned and added into sterile conical flasks each containing 150 ml of medium-enriched seawater. Then, the freshly prepared phytohormones were added to the flasks at specific concentrations (**Supplemental Table S1**). The concentration of phytohormones was based on previous studies (Chu et al. 2019; Fierli et al. 2022; Mc Gee et al. 2020). Each treatment employed three biological replicates. Cultures were cultivated in conical flasks supplemented with various phytohormones, maintaining consistent conditions of light and temperature as those used for stock culture maintenance. The culture medium without the addition of phytohormones was used as the control.

### Experimental data collection

The growth rate of microalgae was monitored by measuring the optical density (OD) of the algal suspension culture every alternate day using a UV-Vis spectrophotometer (Agilent Cary 3500 Multicell) at 680 nm (Paw et al. 2029). For the spectrometric-based quantification of fucoxanthin yield, the absorbance of the cultures was measured at 750 nm on alternate days. Simultaneously, 1 ml of sample from each flask was centrifuged at 7000 rpm and the pellet was resuspended in 1 ml of ethanol. The absorbance of the supernatant was then measured at 445 and 663 nm within 5 minutes of extraction (Wang et al. 2018) and the fucoxanthin yield in cultures supplemented with phytohormones was calculated (**Equation 1**).

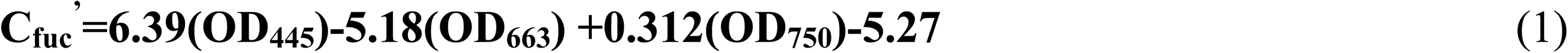

where OD_445_, OD_663,_ and OD_750_ are the absorbance at 445 nm, 663 nm, and 750 nm respectively.

For the estimation of fresh weight biomass, pre-weighed Eppendorf loaded with 1 ml of harvested sample was weighed and was allowed to dry at 60°C till constant weight was obtained to determine the dry weight biomass. The fresh weight and dry weight were calculated by taking the difference between the initial and final weight. This experiment was conducted for 30 days by measuring the growth rate, biomass, and fucoxanthin yield on alternate days.

### Statistical data analysis

We have performed exploratory data analysis to understand the pattern of data and the correlation between the input parameters. Therefore, this study evaluates the efficacy of statistical tools and four ML models in predicting the yield of fucoxanthin in *Isochrysis galbana* using the experimental data as input. All further statistical modelling and figure generation were performed using JMP^®^ (Version 17) and R (R Core Team, 2017).

## Morphological data acquisition

### FESEM data extraction

The morphological changes in the structure of *Isochrysis galbana* without hormone treatment (control) and cultures treated with varying concentrations of hormones were assessed by field emission scanning electron microscopy (FESEM, Thermo Fisher FEI QUANTA 250 FEG).

### Confocal data extraction

Confocal laser scanning microscopy was employed to scrutinize the fluorescence of chlorophyll, lipids, and pigments within the cells. The cell suspensions of *Isochrysis galbana* without hormone treatment (control) and cultures treated with hormones were harvested and subsequently centrifuged at 8000 rpm for 5 min. Pellets were resuspended in PBS buffer (Yadav et al. 2023a). Nile red (9-diethylamino-5Hbenzo [α] phenoxazine-5-one) staining was performed 15 min before imaging to detect the presence of lipid by adding 380 µL of microalgal suspension to 20 µL of Nile red (Sigma; stock solution of 0.2 mg/mL in DMSO). Around 5 µL of the algal suspension was loaded and images were recorded using Confocal Laser Scanning Microscope-Fluoview Fv3000 at 40x objective (Olympus, Japan). The detection ranges were as follows: λ_exc_=488 nm and λ_em_=510-630 nm for carotenoids, λ_exc_=560 nm and λ_em_=640-750 nm for chlorophyll, and λ_exc_=530 nm and λ_em_=636 nm for nile red (Zienkiewicz et al., 2020; Duval et al., 2023).

### Modelling methods

This study utilized four models (RF, SVM, LR, and ANN) to predict the optimized concentration and type of hormone for enhanced fucoxanthin productivity. The performance of these models was compared for the selection of the optimal prediction model. These models are chosen owing to their ability to analyze complex biological data (Kang et al. 2023).

RF is one of the most used ML-based ensemble-learning methods which constructs a forest using multiple decision trees for training and predicting the samples by random extraction (Chen et al. 2023). Each decision tree generates the identification output for unknown test data. Based on the identification output of all decision trees, the final identification output is generated for the unknown test sample. The greater the number of output times for a specific category, it is more likely that the unknown test data belong to it. The process of calculation is simple and easy to understand and interpret, yet it could lead to overfitting performance. The parameters employed for computation of output by RF include the number of trees as 10 and the number of attributes considered at each spit as 6. The features utilized for RF are replicable training and the number of features in the subset could not be less than 4.

SVM is one of the supervised learning methods of ML algorithms that work based on statistical learning theory (Pisner and Schnyer 2020). It is effective in high-dimensional space and could be used for identification, regression, and classification tasks and could function better in conditions where the number of dimensions is higher than the number of samples. The data is effectively separated between two categories using a hyperplane for 2-dimensional data followed by mapping of test points and prediction of its category depending on the side of the gap they belong to. This method could solve the computational complexity and high dimensional issues efficiently. The major disadvantage is it has less sensitivity to data and hence it is strenuous to find appropriate kernel functions for non-linear data. In this study, the parameters for SVM were given as cost (c) is 1, Regression loss epsilon (ε)=0.10, Tolerance limit=0.0010. The radial basis function (RBF) kernel was employed in this study and the iteration limit was set to 100.

LR falls within the realm of supervised machine learning algorithms which operate by learning from labelled datasets and fitting the data points to optimal linear functions. These functions can then be used to predict outcomes for new datasets. It is effective in predictive analysis and provides a linear relationship between dependent and independent variables for the prediction of outcomes (Maulud and Abdulazeez 2020). The Lasso regression (L^1^ norm) was utilized for linear regression with a regularization strength of α=0.001.

ANN could train itself for the recognition of patterns in a dataset and the prediction of non-linear relationships between input variables and output (Kumar et al. 2024). It is demonstrated to be the research hotspot in the field of artificial intelligence and is commonly referred to as a neural network (Chen et al. 2023). A multilayer fully connected feed-forward ANN was applied in this study to develop a model for the prediction of fucoxanthin yield (**Supplemental Fig. S1**). It comprises an input layer, an output layer, and one or more hidden layers. Although the flexibility of the model could be enhanced by increasing the number of hidden layers, one hidden layer is adequate to model the microalgal growth. The process was repeated until the MSE (mean squared error) was achieved as low as possible. All the ML models and the data processing process of this study have been developed using the JMP and Orange software.

### Construction of models for prediction of fucoxanthin yield

The ML models thus developed were used as the driving engine and compared for the accuracy of prediction based on the data used to train the model. In this study, two ML frameworks (Case Study 1 and Case Study 2) were constructed for the inclusion and exclusion of hormone descriptors to train the model and compare its prediction accuracy.

### Case Study 1 (without descriptors)

The ML framework is constructed in a way that when the concentration of hormones, number of days, growth rate, dry biomass, and fresh biomass are given as input, the model will be able to predict the fucoxanthin yield as output. For the initial model, no descriptors will be given for the hormones, hence the prediction will be completely based on the input parameters.

### Case Study 2 (with descriptors)

We have constructed and developed an integrated ML framework to incorporate the characteristics of hormones, hence descriptor values were given to the hormones in addition to the pre-processed experimental data. The input data including days, concentration, and descriptors of hormones will be given as input to the first model which will predict the growth rate. The output of the first model (i.e.) predicted growth rate will be given as input to the second model which will finally predict the fucoxanthin yield.

## Data evaluation

### Evaluation of model performance

To evaluate the accuracy of ML models, 70% of the sample data were selected as the training data set and the remaining 30% were used as the testing dataset. The ML models are trained with the experimental data obtained from supplementation of IAA, SA, GA_3,_ and MeJa phytohormones whereas abscisic acid hormone is used as testing data. The modeling process was repeated 200 times to minimize the errors. The prediction accuracy of the ML models was evaluated using 4 indicators: the coefficient of determination (R^2^), Root Mean Squared Errors (RMSE), Mean Squared Error (MSE), and Mean Absolute Error (MAE) (**Equations 2, 3, 4, 5**) respectively. Therefore, these indicators could better measure the degree of fitness between actual and simulated values.

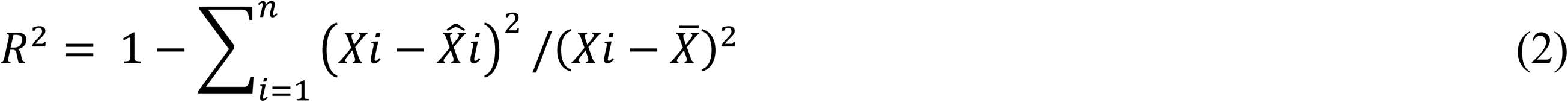

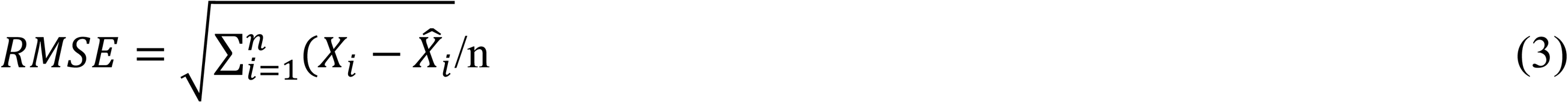

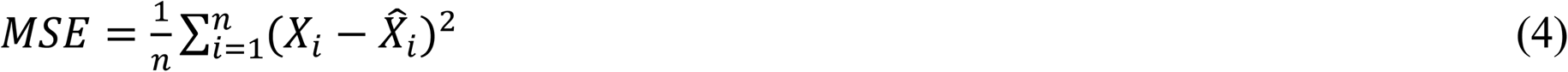

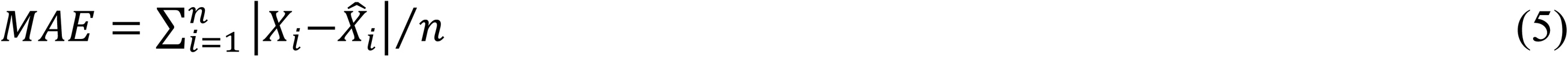

where n is the total number of samples, *X*_i_ and *X̂*_*i*_ are the actual measured and predicted fucoxanthin yield of the samples, respectively, and *X̄* denotes the mean of the measured fucoxanthin yield.

## Results

### Data acquisition and visualization

The proposed approach endorsed simultaneous data collection from the *Isochrysis galbana* to monitor the growth as well as cellular production of fucoxanthin and biomass. The spectrophotometric quantification of fucoxanthin showed advantages in terms of time and cost. The default experimental setup ensured that the result of the proposed method would not be affected by temperature and light. Hence the results will be impacted by the type and concentration of hormones and number of days. I 1, I 2, I 3, and I 4 indicate the hormones IAA, SA, GA_3,_ and MeJa respectively.

### Statistical models and correlation analysis

The data from the supplementation of four phytohormones were analyzed by statistical analysis (scatter plot) to explore and understand the data distribution across various parameters (**Fig. 2**). Among them, the growth rate of microalgae is directly proportional to the fucoxanthin yield. Furthermore, the days at which maximum growth rate and fucoxanthin yield were attained are tabulated (**Supplemental Table S2)**. Overall, the maximum fucoxanthin yield was achieved with 0.2 mg/L MeJa supplementation in minimal time within 10 days.

**Figure 2.**
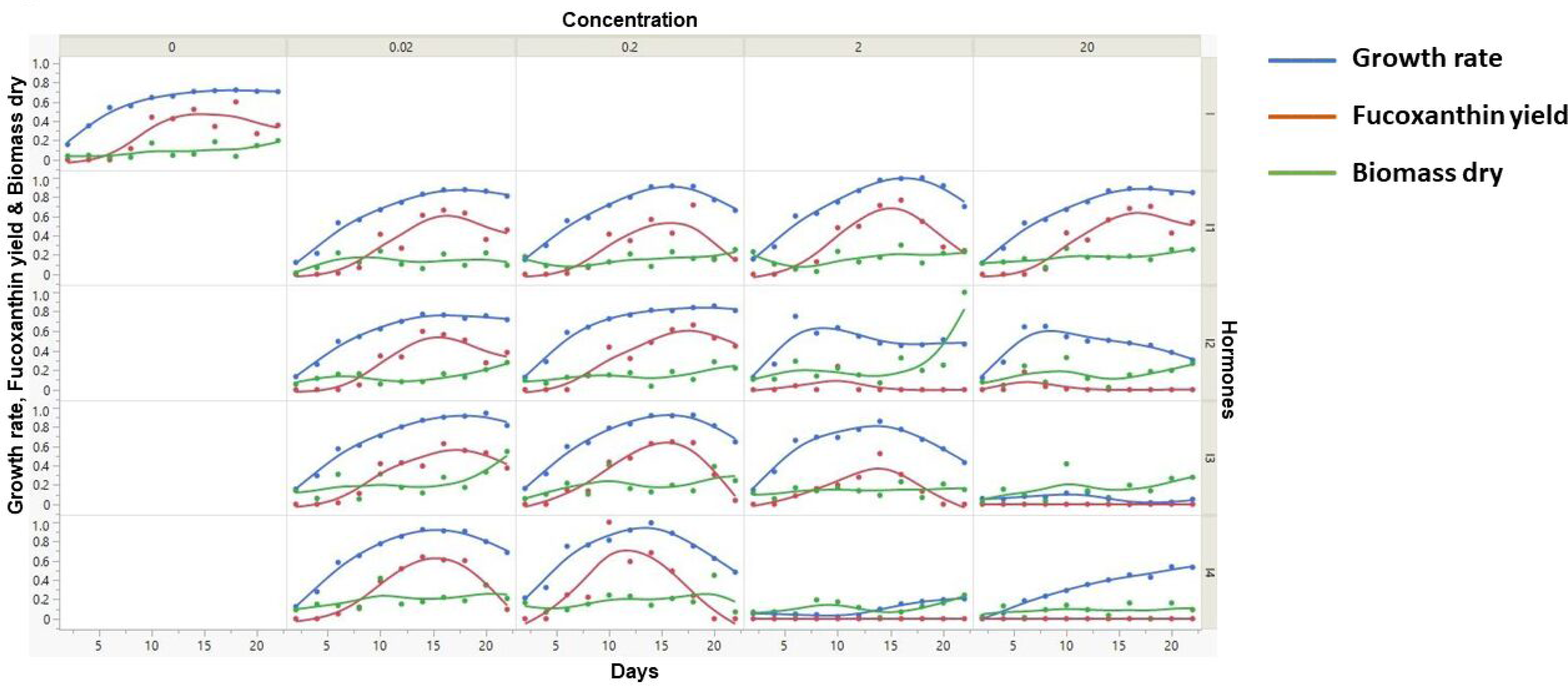
Scatter plot analysis of various input parameters against fucoxanthin yield. x-axis represents the number of days while y-axis (left) represents the fucoxanthin yield. Furthermore the x and y axes are further partitioned into five partitions representing the type and concentration of hormones.

In this study, the experimental data excluding hormone descriptors and including hormone descriptors allowed Pearson correlation coefficient analysis of fucoxanthin across the investigated complete set of input parameters respectively (**Fig. 3**). On a relative basis, consistent with the scatter plot analysis, the fucoxanthin yield showed maximum correlation against growth rate followed by dry weight of biomass whereas concentration and number of days show a negative correlation in both cases (**Fig. 3A and 3B**), demonstrating a weak and moderate association. When descriptors of hormones were included in the input data (**Fig. 3B**), the fucoxanthin yield showed a minimal positive correlation with the hydrogen bond donor count depicting a moderate association.

**Figure 3.**
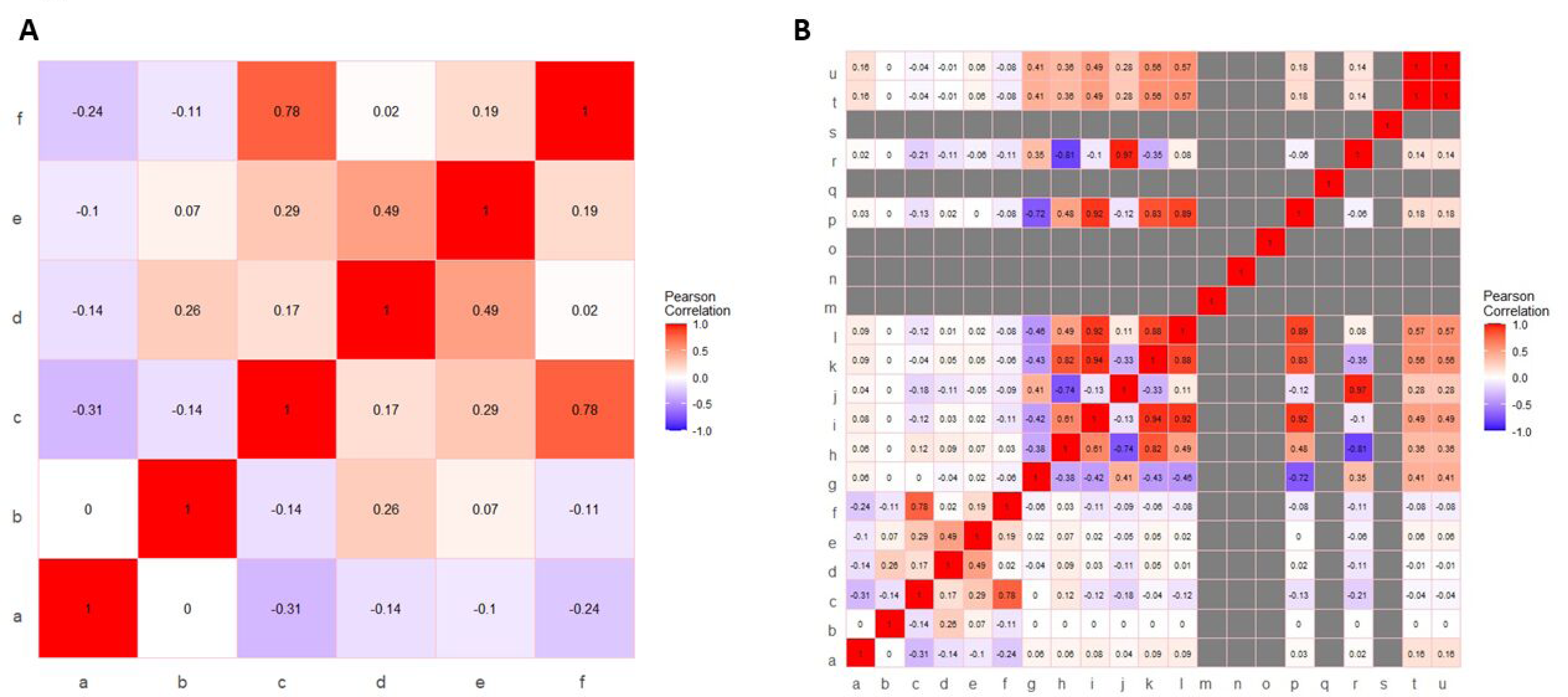
Pearson correlation analysis of input variables (excluding and including) hormone descriptors. **Fig. 3A. Pearson correlation analysis of input variable excluding hormone descriptors** (Notations for figure, a-Phytohormone concentration; b-Days; c-Growth Rate; d-Biomass (Wet), e-Biomass (Dry), f-Fucoxanthin Yield). **Fig. 3B. Pearson correlation analysis of input variable including hormone descriptors** (Notations for figure, a-Phytohormone concentration; b-Days; c-Growth Rate; d-Biomass (Wet), e-Biomass (Dry), f-Fucoxanthin Yield, g-XLogP3; h-Hydrogen Bond Donor Count; i, Hydrogen Bond Acceptor; j, Rotatable Bond Count; k, Topological Polar Surface Area; l, Heavy Atom Count; m, Formal Charge; n, Complexity; o, Isotope Atom Count; p, Defined Atom Stereocenter Count; q, Undefined Atom Stereocenter Count; r, Covalently-Bonded Unit Count; s, Defined Bond Stereocenter Count; t, Undefined Bond Stereocenter Count; u, Canonicalized compound).

### Morphological alterations in microalgae structure

FESEM analysis revealed the morphology of *Isochrysis galbana* cells at day 12 in the absence of hormone treatment and at various concentrations of hormone treatment (**Fig. 4)**. At control, the cells appear clustered with smooth surfaces whereas different concentrations of hormone treatment morphologically alter the structure of microalgae. The caption of each figure indicates the average diameter of the microalgal cell followed by the yield of fucoxanthin

(**Fig. 4**). For instance, IAA when supplemented at 0.02 and 0.2 mg/L, the cells appear clustered and enlarged whereas higher concentrations caused the cell surface to become relatively rough with irregular grooves and increase the average cell size. In contrast, SA (0.02 and 0.2 mg/L) concentrations depict enlarged cells with rough and distorted cell surfaces. SA (2 and 20 mg/L) makes the cells appear smaller with no proper shape. The cells appear smooth round and enlarged at 0.02 mg/L concentration of GA_3_ whereas at 0.2 and 2 mg/L concentrations, they become irregularly shaped and stretchy respectively. The supplementation of GA_3_ at 20 mg/L clusters the cells with irregular grooves and protuberances on the surface. On the other hand, cells appear smooth and round at MeJa (0.02 mg/L) and swollen and enlarged with maximum production of fucoxanthin at MeJa (0.2 mg/L). MeJa (2 mg/L) supplementation completely alters the cell with a distorted and irregular shape. At 20 mg/L the cells appear extremely swollen which causes the cell to explode. Hence, the morphology of cells was altered depending on the type and concentration of the hormone supplementation to medium.

**Figure 4.**
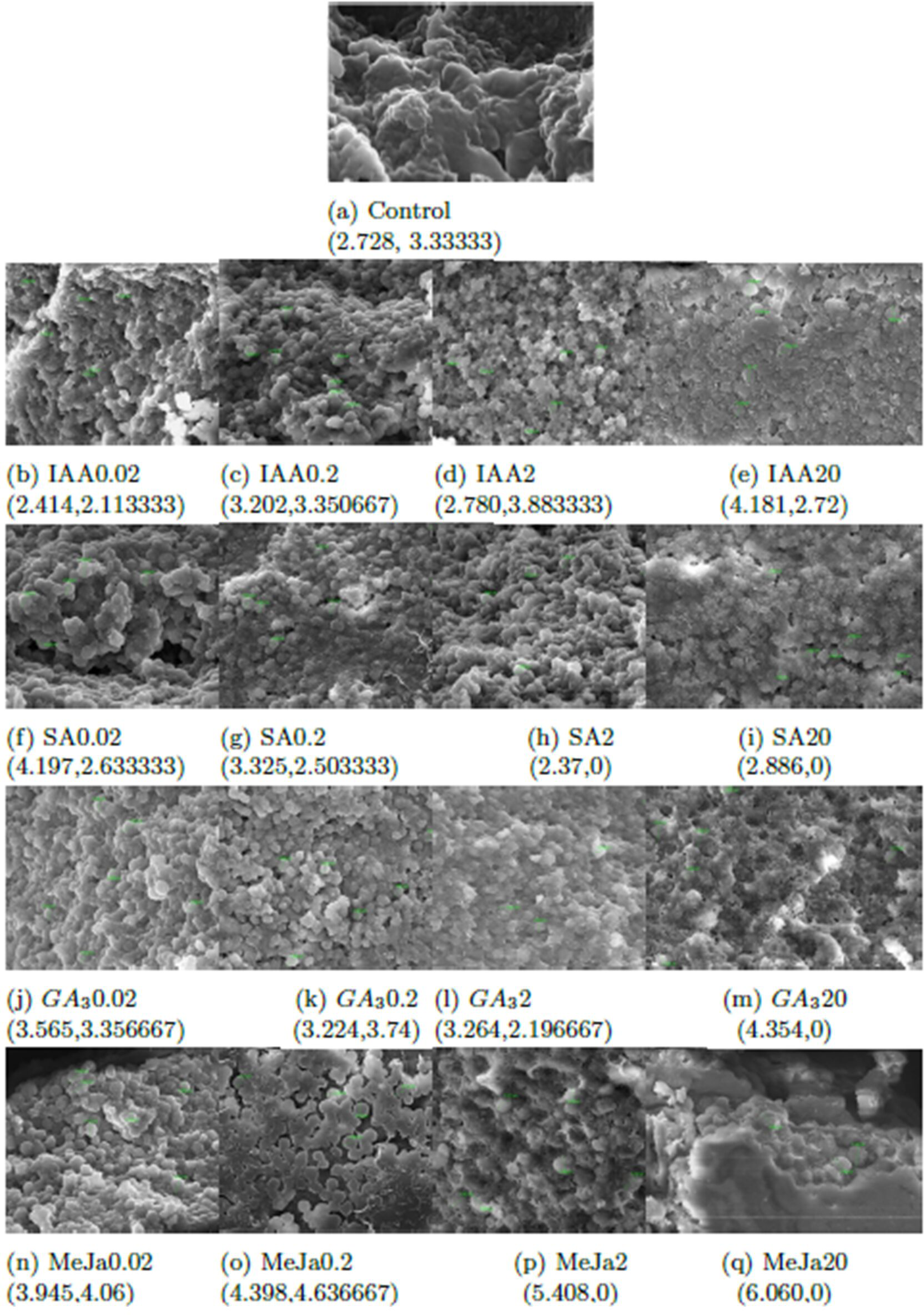
**FESEM analysis of *Isochrysis galbana* without and with hormone treatment.** The caption of each hormone concentration indicates the average cell size and the yield of fucoxanthin.

### Confocal microscopic analysis

In this study, we visualized the impact of hormone treatment on the structure and presence of lipid, pigment, and chlorophyll within *Isochrysis galbana*. Carotenoids are lipophilic pigments present in the interior and exterior of chloroplasts and are detected as green globular forms using Nile red stain under confocal microscopy whereas chlorophyll auto-fluoresce as red globules. The chlorophyll autofluorescence of *Isochrysis galbana* cells in the exponential phase reveals that the type and concentration of phytohormone supplementation negatively affects the chlorophyll content. For instance, supplementation of IAA and SA at higher concentrations demonstrated higher chlorophyll content whereas supplementation of GA_3_ and MeJa at higher concentrations demonstrated degradation of chlorophyll (**Fig. 5A**). During IAA and SA supplementation, the lipid droplets increased in size and number whereas GA_3_ and MeJa supplementation progressively decreased the size of lipids (**Fig. 5B**). As fucoxanthin belongs to xanthophyll carotenoids, the carotenoid fluorescence emission is detected as green globules at 488 nm excitation whereas chlorophyll was detected by red light excitation at 560 nm. Similar to lipids, the type and concentration of hormone supplementation affected the accumulation of carotenoids. MeJa (0.2 mg/L) showed the maximum carotenoid accumulation (**Fig. 5C**). The merged fluorescence emitted by lipids and pigment and chlorophyll autofluorescence within *Isochrysis galbana* in the absence and presence of varying concentrations of hormone supplementation (**Fig. 5D**).

**Figure 5.**
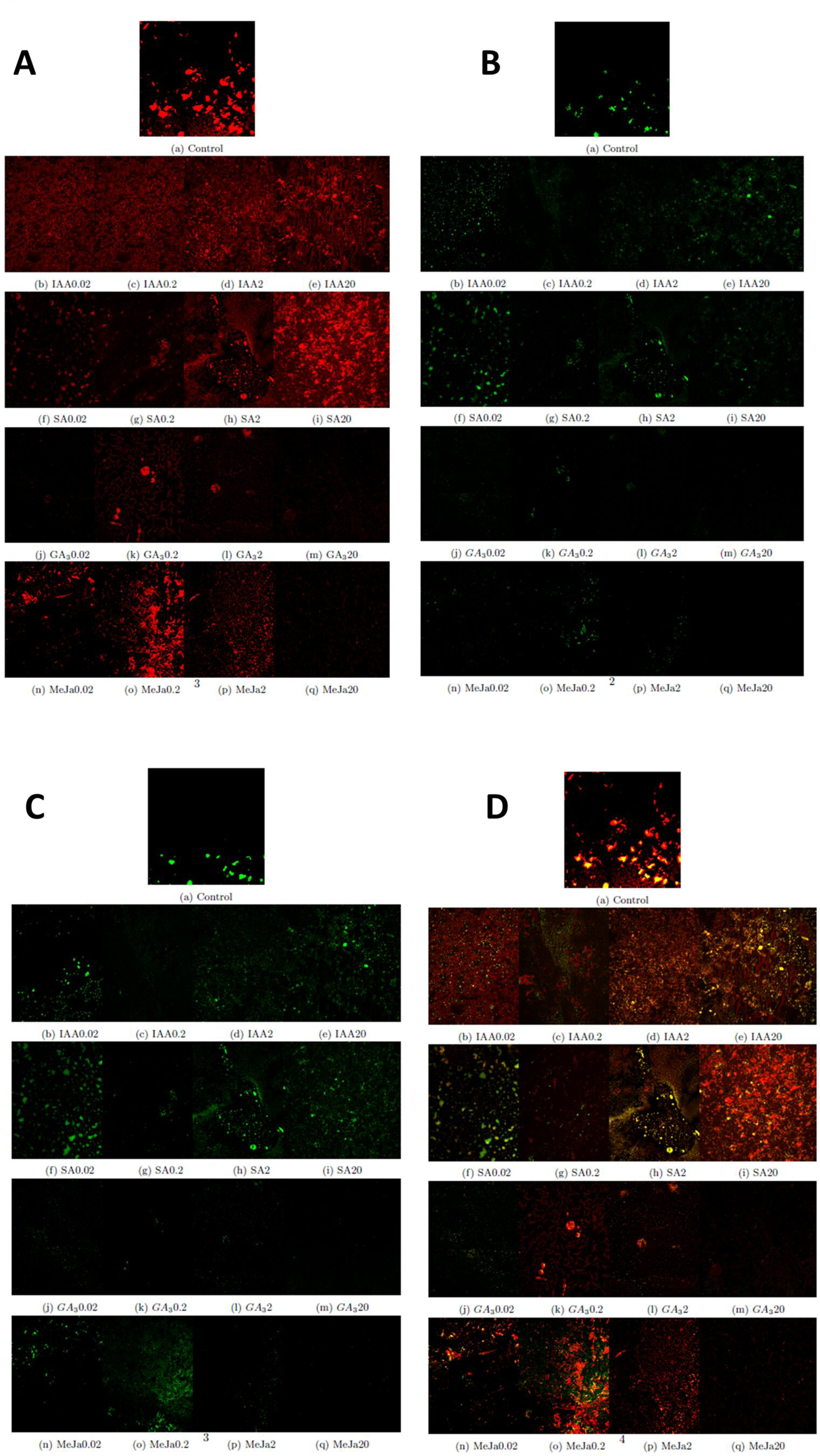
Cellular changes in chlorophyll, lipid, and pigment content of *Isochrysis galbana* in response to phytohormone supplementation. (Bars= 10 µm). **Fig. 5A.** Chlorophyll autofluorescence (red) of *Isochrysis galbana* cells in response to phytohormone supplementation. **Fig. 5B**. Nile red stained Lipid fluorescence (green) of *Isochrysis galbana* cells in response to phytohormone supplementation. **Fig. 5C**. Pigment fluorescence (green) of *Isochrysis galbana* cells in response to phytohormone supplementation. **Fig. 5D**. Merged fluorescence of chlorophyll, lipid and pigment of *Isochrysis galbana* cells in response to phytohormone supplementation.

### Fucoxanthin prediction using different models

In this study, the experimental dataset was divided into training and testing data and four ML models (SVM, RF, LR, ANN) were adopted for the prediction of fucoxanthin yield as these models are extensively used for analyzing complex biological data (**Fig. 6A**). The reliability of the models was evaluated based on previously trained data. The parameters considered for the construction of ML models and framework of models for Case Study 1 and Case Study 2 are illustrated in (**Fig. 6B**). For Case Study 1, the models were trained using whole experimental data whereas for Case Study 2, models were trained with restricted experimental data, and descriptors for hormones were included to incorporate the characteristics of hormones. The experimental dataset (growth rate, dry biomass, fresh biomass, number of days, type and concentration of hormones IAA, SA, GA_3_ MeJa) used for the modelling step in this study included a dataset consisting of 273 samples with 6 variables (excluding hormone descriptors) for the Case Study 1, whereas, for Case Study 2, the experimental dataset consisted of 273 samples with 24 features (including hormone descriptors) for prediction of fucoxanthin yield.

**Figure 6.**
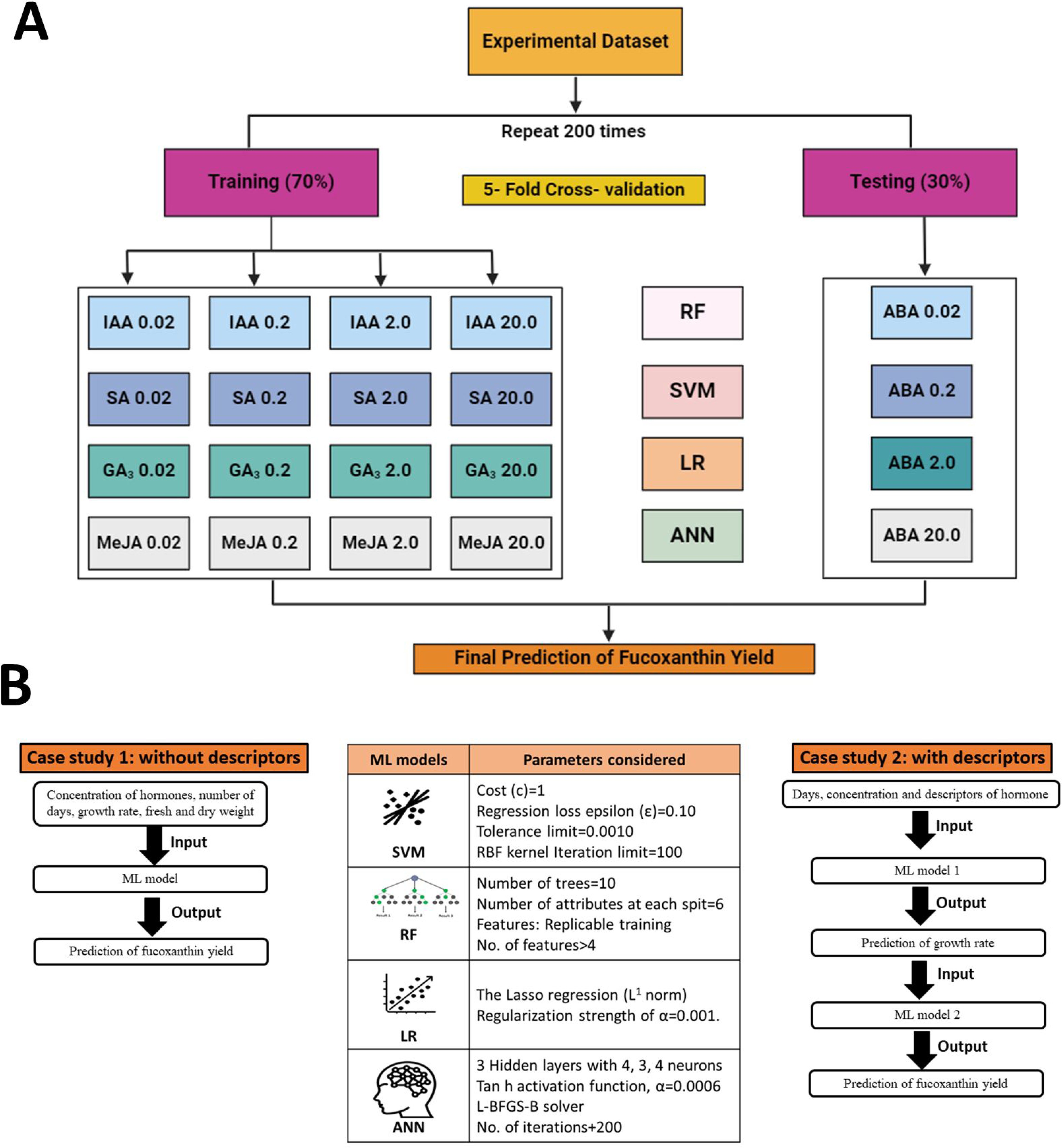
Overall overview of ML framework construction and training and testing dataset. **Fig. 6A.** Training and testing dataset used to train ML models **Fig. 6B**. Construction of ML frameworks for Case study 1 and Case study 2

### Case Study 1 (excluding hormone descriptors)

For Case Study 1, the RF model performance for fucoxanthin yield prediction provided higher *R*^2^ values and lower RMSE, MSE, and MAE values (*R*^2^ = 0.809, *MSE* = 0.602, *RMSE* = 0.776, *MAE* = 0.458). For the prediction of fucoxanthin yield (**Table 1**), *R*^2^ values of LR and SVM were lower than the RF and ANN models. Among the 4 models, RF provided the maximum accuracy in fucoxanthin yield prediction followed by the ANN model (*R*^2^ = 0.722). Compared with other models, LR was the poorest algorithm for predicting the fucoxanthin yield (*R*^2^ = 0.605, *MSE* = 1.248, *RMSE* = 1.117, *MAE* = 0.906). The randomly selected predictions made by four ML algorithms at specified instances show that RF and ANN models predicted the fucoxanthin yield with maximum accuracy and suggested that MeJa (0.2 mg/L) proved to synthesize maximum fucoxanthin compared to other hormones (**Table 1**). Therefore, RF and ANN models were adopted as the optimized modeling method for fucoxanthin prediction for Case Study 1.

**Table 1.** Test results of ML models trained with all input variables excluding hormone descriptors.

| Model | MSE | RMSE | MAE | R<sup>2</sup> |
| --- | --- | --- | --- | --- |
| Random Forest | 0.602 | 0.776 | 0.458 | 0.809 |
| Linear Regression | 1.248 | 1.117 | 0.906 | 0.605 |
| Support Vector Machine | 0.949 | 0.974 | 0.648 | 0.699 |
| Artificial Neural Network | 0.878 | 0.937 | 0.506 | 0.722 |

The randomly selected ML-predicted fucoxanthin yield at different days, types, and concentrations of hormones show the differences in the prediction of ML models (**Supplemental Table S3**). From these predictions, it can be inferred that MeJa could yield maximum fucoxanthin production at lower concentrations in a shorter time followed by GA_3_. The hormones IAA and SA were able to produce higher fucoxanthin after 15 days. However, the results obtained by ML prediction were purely based on the training and experimental data as the hormone descriptors have not been included. Hence, descriptors of hormones were given as an additional input to the developed model and prediction performance was evaluated (**Supplemental Table S4**). Consistent with the previous results, RF and ANN models (*R*^2^ = 0.825, *R*^2^ = 0.746) showed better predictions respectively. This result suggests that the inclusion of hormone descriptors in input data improved the prediction accuracy of fucoxanthin yield (**Supplemental Table S5**).

### Case Study 2 (including hormone descriptors)

#### Growth rate prediction using pre-processed data

As the inclusion of hormone descriptors improved the prediction accuracy (Case Study 1), a generic integrated ML model was constructed exclusively to incorporate the hormone characteristics and the predictive performance of ML models for growth rate and fucoxanthin yield has been evaluated. The experimental data was pre-processed before training the ML models to avoid discrepancies. In this model, growth rate and fucoxanthin yield will be predicted by varying the input data (**Fig. 6B**) The prediction results of ML models (**Table 2**) showed ANN to predict growth rate with maximum accuracy (*R*^2^ = 0.836) followed by RF (*R*^2^ = 0.82) respectively. These results indicate that the ANN model showed better performance in the prediction of the growth rate in several instances whereas LR showed the poorest prediction accuracy of the growth rate. However, RF failed to provide the expected maximum estimated accuracy at growth rate prediction, which was provided by ANN (**Supplemental Table S6**). These results demonstrate that the ANN model performed better than the other models in predicting the growth rate of *Isochrysis galbana*.

**Table 2.** Test results of ML models for the prediction of growth rate using pre-processed data.

| Model | MSE | RMSE | MAE | R<sup>2</sup> |
| --- | --- | --- | --- | --- |
| Random Forest | 0.011 | 0.105 | 0.083 | 0.820 |
| Linear Regression | 0.052 | 0.227 | 0.195 | 0.157 |
| Support Vector Machine | 0.021 | 0.144 | 0.116 | 0.661 |
| Artificial Neural Network | 0.01 | 0.1 | 0.077 | 0.836 |

#### Fucoxanthin prediction using pre-processed data

In this study, the ML models were fed with predicted growth rate as input data for the prediction of the fucoxanthin yield by combining the advantages of integration of the ML models and avoiding overfitting or overestimating. Compared to all the above models, the RF model (*R*^2^ = 0.839) employed in this method gave the maximum accuracy for fucoxanthin yield prediction followed by ANN model (*R*^2^ = 0.738) whereas the predictive performance of SVM and LR were better than the previously developed models (**Table 3**). In several instances, the RF model fed with pre-processed experimental data showed a better prediction of fucoxanthin yield followed by the ANN model (**Supplemental Table S7**). For Case Study 2 (including descriptors), the RF and ANN model was able to improve the generalization by integration of multiple models, thus providing a more stable prediction result. The prediction values obtained from the integration of ML models are in good agreement with the measured fucoxanthin yield from *Isochrysis galbana*, which reflects a satisfactory prediction result.

**Table 3.** Test results of generic ML models for the prediction of fucoxanthin yield using pre-processed data.

| Model | MSE | RMSE | MAE | R<sup>2</sup> |
| --- | --- | --- | --- | --- |
| Random Forest | 0.507 | 0.712 | 0.357 | 0.839 |
| Linear Regression | 1.27 | 1.127 | 0.919 | 0.598 |
| Support Vector Machine | 1.332 | 1.154 | 0.781 | 0.578 |
| Neural Network | 0.826 | 0.909 | 0.51 | 0.738 |

#### Prediction of growth rate using raw data

In this study, additionally, to evaluate the influence of pre-processing of experimental data on the prediction of growth rate and fucoxanthin yield, the constructed models were trained with raw data. Consistent with the results for pre-processed data, the growth rate prediction results were better with the ANN model followed by the RF model (**Supplemental Table S8).** ANN model showed the maximum accuracy of growth rate predictions (*R*^2^ = 0.846) whereas LR showed the worst growth rate prediction accuracy (*R*^2^ = 0.156). However, the RF model failed to provide the expected estimated accuracy at growth rate prediction, which was provided by ANN (**Supplemental Table S9**). Hence, the artificial neural network is optimized as the best model for the prediction of growth rate for both raw and pre-processed data. Although the RF model failed to provide the best prediction accuracy for all the case studies, it achieved a more stable performance by minimizing the deviations and randomness of the other models.

#### Prediction of fucoxanthin yield using raw data

In this study, when raw data is given as input, the ANN (*R*^2^ = 0.836) and RF models (*R*^2^ = 0.845) achieved the maximum accuracy in the prediction of fucoxanthin yield. The predictive performance of LR remained the same as that of the model trained with pre-processed data whereas the predictive performance of SVM decreased **(Supplemental Table S10)**. The RF model showed better prediction similar to the measured fucoxanthin yield at several instances whereas ANN overestimated the fucoxanthin yield at few instances (**Supplemental Table S11**). Hence, these results infer that pre-processing of data shows an influence on the predictive performance of the ML models. However, contrary to the ML-based fucoxanthin prediction, the quantitative experimental values of fucoxanthin yield obtained for the *Isochrysis galbana* are relatively lower than that obtained from ML prediction at few instances. This discrepancy is possibly observed as ML models at few instances over-estimated the fucoxanthin production depending on the training dataset.

## Discussion

Microalgae synthesize a wide range of bioactive metabolites including carotenoids, lipids, and polysaccharides which makes them a sustainable source for next-generation feedstock (Foo et al., 2017). The microalgal species *Isochrysis galbana* was selected in this study because they have gained widespread application in aquaculture and animal feed due to their rapid and stable growth rates. However, compared with other microalgal species (*Phaedotactylum tricornutum*, *Chaetoceros calcitrans*), there have been relatively fewer studies on *Isochrysis galbana* for fucoxanthin production. Therefore, the current study, which focuses on predicting the fucoxanthin production of *Isochrysis galbana* through UV spectroscopic method coupled with high-throughput ML studies in the research field, is of great significance for the future development of commercial production of microalgal fucoxanthin. The integration of ML models with biotechnological tools (UV spectrometric based measurement of fucoxanthin yield) allows for the rapid and accurate prediction of fucoxanthin yield which can aid in understanding the influence of different type and concentration of hormones on the microalgal growth, biomass and response to elicitor supplementation. By predicting the fucoxanthin production of *Isochrysis galbana*, this study can provide valuable insights into their enhanced yield potential and optimized type and concentration of hormones, aiding in the improved cultivation strategies as well as commercial fucoxanthin production strategies.

**Figure 1** represents the experimental workflow of UV-based measurement of fucoxanthin coupled with ML-based fucoxanthin prediction whereas **Figure 2** represents the scatter plot analysis of experimental data which shows the type and concentration of hormone supplementation has an influence on the fucoxanthin yield. For control cultures, the maximum yield of fucoxanthin was achieved only on day 18 whereas 0.02 concentration of I 1 and I 2 showed maximum fucoxanthin yield on days 10 to 15. At I 1 (0.2, 2, and 2 mg/L), I 2

(0.2 mg/L), I 3 (0.02 and 0.2 mg/L), maximum yield was obtained from days 15 to 20. The supplementation of I 4 (0.02 mg/L) could give maximum fucoxanthin yield at days 12 to 18 whereas I 4 (0.2 mg/L) concentration could attain maximum yield within 10 days. For hormones I 2, I 3, and I 4 (2, 20 mg/L), there is negligible or minimum yield of fucoxanthin.

These results were consistent with the previous findings on phytohormone supplementation. Mc Gee et al. (2020) showed that fucoxanthin content in *Stauroneis* sp. and *Phaeothamnion* sp. increased owing to the addition of MeJa (10 and 100 µM). It was also reported that MeJa (2.2 mg/L) supplementation enhanced the biosynthesis of fucoxanthin in *Stauroneis* sp. (Mc Gee et al. 2021). Similar results were obtained for *P. tricornutum* cultivated with GA_3_. The supplementation of SA also boosted the synthesis of carotenoids in *Nitzschia* leading to a 1.7-fold increase in fucoxanthin content. In contrast, MeJa supplementation at 0.5 mg/L has a negligible impact on fucoxanthin yield (Fierli et al. 2022).

Additionally, Fierli et al. (2023) studied the effect of the combined application of exogenous phytohormones along with blue light in *Phaeodactylum tricornutum*. GA_3_ when supplemented separately increased the fucoxanthin yield by 30%. The combined supplementation of GA_3_ and ABA was demonstrated to be more effective. Therefore, supplementation of phytohormones provides a promising strategy to enhance fucoxanthin production due to their intrinsic role in promoting microalgal growth. A similar pattern of results was obtained in studies predicting the fucoxanthin production. Gao et al. (2021), studied the effect of light on biomass and fucoxanthin production in *Phaeodactylum tricornutum* and *Tisochrysis lutea*. The prediction models developed using fluorescence spectroscopy showed a positive correlation between biomass and fucoxanthin yield (Gao et al. 2021). However, the impact on fucoxanthin production depends on the type of hormone, concentration, and the microalgal species.

### Pearson correlation coefficient analysis

Pearson correlation coefficient uses a correlation coefficient (*R*) ranging between *-1* to *+1* to evaluate the linear relationship between the variables X and Y. The ideal positive and negative relationship between the variables is indicated by *R* values of *1* and *-1* respectively. The absolute magnitude of R represents the strength of correlation such that a higher absolute value indicates a greater correlation. An absolute value of *R>0.6* is considered a robust correlation. We detected a positive correlation between growth and fucoxanthin yield in both the cases of exclusion and inclusion of hormone descriptors in input data (**Fig. 3A** and **Fig. 3B**) respectively. In this study, the growth rate of *Isochrysis galbana* shows a higher positive correlation with fucoxanthin yield (*R=0.78*). This is consistent with previous studies showing a positive correlation of fucoxanthin yield with microalgal growth rate followed by biomass (Li et al. 2020; Ishika et al. 2019). Recently, Sequeira et al. (2021) reviewed the positive influence of hydrogen bond donor chemicals on the yield of fucoxanthin from macroalgae as well as microalgae. Hence, consistent with theoretical expectations and prior observations, when hormone descriptors are included in input, fucoxanthin yield shows a higher positive correlation with growth rate followed by hydrogen bond donor count.

### Morphological variations in response to hormone treatment

In this study, FESEM analysis of *Isochrysis galbana* at day 12 revealed that the type and concentration of hormones alter the morphological structure of microalgal cells (**Fig. 4)**. Consistent with this result, similar changes in the morphology of microalgae were observed when the concentration of nutrient supplementation was varied. For instance, variations in the nutrient composition of the culture medium morphologically altered the cell wall and structure of *Amphiprora* sp. (Jayakumar et al. 2021).

### Confocal analysis

The presence of lipid, pigment, and chlorophyll within *Isochrysis galbana* was visualized using confocal microscopic analysis. The chlorophyll autofluorescence-based detection method has revealed immense potential as an on-site tool to assess microalgal vitality (Li et al. 2022). However, there is very limited data on the impact of phytohormones on the presence of chlorophyll within the microalgal structure. In this study, the effects of different phytohormones with four concentrations (0.02, 0.2, 2, and 20 mg/L) on the chlorophyll autofluorescence in cells of *Isochrysis galbana* were investigated by red light excitation at 560 nm (**Fig. 5A**). Experimental results showed that both type and concentration of hormones were major factors that cause the degradation of chlorophyll.

There are several reports on the enhanced accumulation of lipids in Nile red-stained microalgal cells grown under nutrient-stress conditions and phytohormone supplementation. In this study, *Isochrysis galbana* cells the lipid accumulation was comparatively higher within the cells supplemented with IAA and SA hormones whereas supplementation of GA_3_ and MeJa at higher concentrations degraded the lipids (**Fig. 5B**). These results were consistent with the previous studies (Ahamed et al. 2022; Duval et al. 2023; Zienkewwicz et al. 2020).

Additionally, spectral analysis using confocal laser scanning microscope was performed to investigate the alterations in fluorescence emission of endogenous pigments in *Isochrysis galbana* cells. The microalgal pigments when excited by specific wavelengths of UV-visible laser light will produce a specific emission spectrum. The fluorescence emission of carotenoids is detected in the green-yellow spectral region whereas chlorophyll is typically detected in the red spectral region (Zienkiewicz et al. 2020). In this study, when blue light at 488 nm excitation was given to hormone-treated *Isochrysis galbana* cells, a change in spectral characteristic occurred owing to an increased carotenoid pigment (**Fig. 5C**).

### Performance of machine learning models for fucoxanthin prediction

Few previous studies have demonstrated the feasibility of the UV spectroscopic method and the fusion of ML models to analyze the data from multiple treatment parameters could provide better prediction of chlorophyll and other pigments. However, fucoxanthin prediction based on the UV-spectroscopic method have not been previously investigated.

### Differences in fucoxanthin prediction metrics (Case Study 1)

It can be observed from the prediction results of ML models trained with whole input data excluding descriptors (**Table 1**), that the RF model is the most stable and showed higher accuracy with less error rate (*R*^2^ = 0.809, *MSE* = 0.602, *RMSE* = 0.776, *MAE* = 0.458) followed by the ANN model (*R*^2^ = 0.722, *MSE* = 0.878, *RMSE* = 0.937, *MAE* = 0.506). Consistent with previous results, ML models trained with whole data including hormone descriptors (**Supplemental Table S4**), the RF and ANN models showed the maximum prediction accuracy. The major advantage of the RF model over the other ML models is that it utilizes an integrated learning algorithm to generate multiple decision trees for learning and prediction. The average of each decision tree was used to attain the final prediction. Thus, this assures robust training and decreases the chances of overfitting and the influence of noised data. On the other hand, LR and SVM models enable single training from the input dataset without statistical average and bootstrap sampling. Hence, compared with other models, RF models show better performance as per previous studies (Chen et al. 2023, Lei et al. 2019).

### Differences in growth rate prediction metrics (Case Study 2)

As in Case Study 1, the inclusion of hormone descriptors in the basic model improved the prediction accuracy of fucoxanthin yield, we constructed an integrated ML model framework exclusively for the inclusion of hormone descriptors and pre-processed the experimental data to avoid further discrepancies. It can be observed that the construction of the ML model (**Fig. 6**) and the inclusion of hormone descriptors in pre-processed input data enhanced the prediction accuracy compared to Case Study 1 (**Table 2 and Supplemental Table S6**). But ANN model showed maximum accuracy in the prediction of the growth rate (*R*^2^ = 0.836) whereas the RF model showed maximum accuracy in the prediction of fucoxanthin yield using pre-processed data (*R*^2^ = 0.839) (**Table 3**). Furthermore, these results are in strong accordance with ML models trained with raw data (**Supplemental Table S8, S9**) as the ANN model showed the maximum growth rate prediction accuracy (*R*^2^ = 0.846).

### Differences in fucoxanthin yield prediction metrics (Case Study 2)

When the constructed model was trained with previously predicted growth rate as input (**Table 3**), the RF model showed the maximum fucoxanthin prediction accuracy (*R*^2^ = 0.839) followed by the ANN model (*R*^2^ = 0.738). The prediction results of fucoxanthin yield by generic integrated ML model trained with growth rate from previous model showed RF to be the best model **(Supplemental Table S7)**.

Consistent with the above results, ML models trained with raw data (**Supplemental Table S10, S11**) gave the best fucoxanthin prediction results with the ANN model (*R*^2^ = 0.836) and RF model (*R*^2^ = 0.845). The predictive performance of LR and SVM were better than the previously developed models. These results are in strong accordance with the concept that neural network effectively processes the non-linear characteristics of data when enough data and neurons are given. For an ANN model to be ideal, it requires three vital functions to operate (Otálora et al. 2021). The major requirement is that the data should be adequate for training and validation of the model. The second vital function is the construction and structure of the neural network which includes the selection of the type, size, and choice of layers based on the problem addressed, input type, amount of data, and complexity of the model to be developed. The final part of developing an ideal model lies in the process of training which is defined by the calculation frequency of input parameters, duration of the training, type of data used for training, and the stop factors (Hudson et al. 1992). On the other hand, as SVM and LR process only the linear characteristics of data, they demonstrated poor performance in both Case Study 1 and Case Study 2.

### Performance of ML models (Test data) in fucoxanthin prediction

Marine biotechnological research is progressing swiftly, with a burgeoning interest in utilizing multi-omics approaches and machine learning techniques to analyze marine metabolite datasets (Manochkumar et al. 2023). The developed integrated ML model harnesses the complementary strengths of the basic models to minimize the occurrence of random errors, thereby enhancing the reliability of its predictions. When Abscisic acid phytohormone (predictions and actual measured values are highlighted in red) was used as testing data, RF and ANN networks showed the maximum prediction accuracy (**Supplemental Tables S3, S5, S6, S7, S9, S11**). Even though RF showed stable prediction, the ML models overestimated the fucoxanthin yield of abscisic acid in several instances when compared to the actual measured values. This requires the need to train the ML model with increased sample size and different phytohormones.

Yadav et al. (2023b), investigated the impact of ANN-GA model and statistical RSM-based model to optimize the process parameters and elevate the production of isoprene in engineered *Synechococcus elongatus* UTEX 2973. ANN-GA model combined with metabolic pathway inhibition strategy performed better than the statistical model and achieved a 29.52-fold higher isoprene yield. Similarly, Kang et al. (2023) demonstrated the ML-guided prediction of engineered *Deinococcus radiodurans* R1 for enhanced lycopene production. The multilayer perceptron models combined with the genetic algorithm predicted the potential overexpression targets from 2047 combinations of key genes. This model achieved a 3-fold increased lycopene production from glycerol and a 6-fold increased lycopene yield. Yeh et al. (2023), investigated the use of ML models for modelling and growth monitoring of microalgae. The performance of Long Short-Term Memory (LSTM) and Support Vector Regression (SVR) was compared for outdoor cultivation of *P. tricornutum* in flat-panel airlift photo-bioreactors. The LSTM model outperformed the SVR model and showed its potential ability to capture the acclimation effects of light on microalgal growth. Recently, data-enhanced interpretable ML was used to predict the biochar characteristics. Data enhancement significantly improved the model accuracy from 5.8% to 15.8%. Compared to ANN and SVM, the optimal RF model showed a maximum accuracy of 94.89% (Chen et al. 2023).

Consistent with previous studies, in our study, the addition of hormone descriptors and pre-processing of data to the constructed generic integrated model enhanced the performance of the RF optimal model to 83.9%. Therefore, the production of fucoxanthin depends on the type and concentration of hormone supplementation and number of days of cultivation. In addition, the growth rate of microalgae was directly proportional to the fucoxanthin production. Machine learning models predicted that supplementation of MeJa (0.02, 0.2 mg/L) contributed to maximum fucoxanthin production in shorter time intervals whereas IAA supplementation showed maximum fucoxanthin production in day 18. The created generic model was found to be more effective in predicting the fucoxanthin yield as this is the first study to employ ML to predict the fucoxanthin yield from microalgae.

Testing the potential combination of phytohormones to forecast the synergetic effect on fucoxanthin production and dynamics of microalgal growth will constitute a significant aspect of the upcoming research endeavours in this field. It will be intriguing to contrast various deep-learning models with the ML models employed in this study for the enhancement of fucoxanthin production. Overall, the fucoxanthin production from *Isochrysis galbana* was validated and verified by the construction of different ML models. These constructed models were only applicable in the determination of fucoxanthin yield using the spectrometric-based data acquisition. Therefore, ML models could be applied as a prediction tool for the commercial production of fucoxanthin by tracking the growth rate as well as determining the fucoxanthin yield for industrial purposes. This approach can aid in saving time, costs, and manpower associated with optimizing the process parameters.

### Future improvements

Although the current results are satisfactory, there are still areas for improvement that should be addressed in future research. To further enhance the prediction accuracy of fucoxanthin using machine learning models, expanding the dataset size could prove beneficial, considering that the sample size utilized in this study has certain limitations. For instance, as we trained the model with limited data using four phytohormone supplementations and four concentrations, the ML model could only capture and program as per the characteristics of the trained hormones. Future studies should include more types and concentrations of hormones to test the applicability and robustness of developed ML models. Recent studies have shown that deep learning models can effectively harness large datasets. Therefore, the incorporation of deep learning should be considered to explore the potential applicability of UV-based fucoxanthin detection in marine research.

## Conclusion

This study explored the potential of integrating UV-based fucoxanthin estimation with ML model prediction to establish a reliable predicting tool and examined how various types and concentrations of hormones affect the accuracy of fucoxanthin prediction. The conclusions are summarized as follows

1) Among the supplemented phytohormones, MeJa (0.2 mg/L) enhanced the fucoxanthin production (7.83 μg/ml) in a shorter time interval of less than 10 days.
2) In the case of fucoxanthin prediction based on whole input data, the difference in estimation accuracy between RF and ANN was not substantial. Although the basic models (LR and SVM) demonstrated some capability to predict fucoxanthin yield, the accuracies and stabilities were poor.
3) Compared with the basic ML models of Case Study 1, the integrated ML model (Case study 2) contributed to higher prediction accuracy in most cases. ANN showed maximum accuracy in the growth rate prediction whereas RF showed maximum accuracy in the fucoxanthin prediction.
4) The fucoxanthin prediction accuracy gradually improved with the pre-processing of data and inclusion of hormone descriptors in input data.

The results reported herein reflect the effectiveness of machine learning and UV-based data acquisition for predicting the fucoxanthin yield. This study provides valuable insights to accelerate and enhance fucoxanthin production and pave the way to different ideas for easy prediction of fucoxanthin yield. However, additional studies should be conducted on a larger number of samples treated with more types and varying concentrations of hormones to test and verify the robustness and adaptability of the proposed method.

## Acknowledgments

The authors thank Dr. P. Santhanam, Department of Marine Science, Bharathidasan University for providing the marine microalgae, *Isochrysis galbana.* We thank the VIT management for the constant support.

## Author Contributions

J.M. Conceptualization of experimental research, data analysis, Writing – original draft; A.J. and A.K.C. performed and validated the ML experiments, review & editing; B.V. and D.J. review & editing; S.R. Conceptualization, Supervision, Validation, review & editing.

## Supplemental data

The following materials are available as supplemental data

**Supplemental Figure S1.** Schematic representation of Artificial neural network ML model

The input layer comprises four neurons corresponding to the four input variables (type of hormone, concentration of hormones, growth rate and biomass) and the output layer has one neuron corresponding to fucoxanthin yield. Three hidden layers with 4, 3, and 4 neurons respectively were found to give good performance. The tan h activation function was utilized in this study for the computation of output. The regularization value of α=0.0006 and L-BFGS-B solver was used for the ANN model. The neurons between adjacent layers are fully interconnected and the number of iterations was set to 200 for the training algorithm to reduce the error between the actual and predicted output.

**Supplemental Table S1.** Concentrations of phytohormones used for culturing *Isochrysis galbana*.

**Supplemental Table S2.** Maximum growth rate and fucoxanthin yield attained for various phytohormones.

**Supplemental Table S3.** Prediction of fucoxanthin yield by ML models trained with all input variables excluding hormone descriptors

**Supplemental Table S4.** Test results of ML models trained with all input variables including hormone descriptors.

**Supplemental Table S5.** Prediction of fucoxanthin yield by ML models trained with all input variables.

**Supplemental Table S6.** Prediction of growth rate by ML models trained with restricted pre-processed input data

**Supplemental Table S7.** Prediction of fucoxanthin yield by generic ML models trained with restricted pre-processed input data

**Supplemental Table S8.** Test results of growth rate prediction by ML models trained with raw data.

**Supplemental Table S9.** Prediction of growth rate using raw data.

**Supplemental Table S10.** Test results of fucoxanthin prediction by ML models trained with raw data.

**Supplemental Table S11.** Prediction of fucoxanthin yield using raw data.

